# Distinct roles for SOX2 and SOX21 in differentiation, distribution and maturation of pulmonary neuroendocrine cells

**DOI:** 10.1101/2022.01.18.476762

**Authors:** Evelien Eenjes, Anne Boerema-de Munck, Marjon Buscop-van Kempen, Dick Tibboel, Robbert J. Rottier

## Abstract

Pulmonary neuroendocrine (NE) cells represent a small population in the airway epithelium, but despite this, hyperplasia of NE cells is associated with several lung diseases, such as congenital diaphragmatic hernia and bronchopulmonary dysplasia. The molecular mechanisms causing the development of NE cell hyperplasia remains poorly understood. Previously, we showed that the SOX21 modulates the SOX2-initiated differentiation of epithelial cells in the airways (Eenjes, 2021). Here, we show that precursor NE cells start to develop in the SOX2+SOX21+ airway region and that SOX21 suppresses the differentiation of airway progenitors to precursor NE cells. During development, clusters of NE cells start to form and NE cells mature by expressing neuropeptide proteins, such as CGRP. Deficiency in SOX2 resulted in decreased clustering, while deficiency in SOX21 increased both the numbers of NE ASCL1+ precursor cells early in development, and the number of cell clusters at E18.5. In addition, at the end of gestation (E18.5), a number of NE cells in *Sox2* heterozygous mice, did not yet express CGRP suggesting a delay in maturation. In conclusion, SOX2 and SOX21 function in the initiation, migration and maturation of NE cells.

## INTRODUCTION

The airway epithelium consists of different cell types, such as basal, secretory and ciliated cells. A rare (<1%) subpopulation of airway epithelial cells, the pulmonary neuroendocrine (NE) cell, is associated with a variety of lung diseases (Garg, Sui et al. 2019). These lung diseases may have a complex origin, showing an inflammatory phenotype like bronchopulmonary dysplasia (BPD), asthma and chronic obstructive pulmonary disease (COPD) (Cutz, Yeger et al. 2007, Gu, Karp et al. 2014, Sui, Wiesner et al. 2018) or may have a congenital origin like congenital diaphragmatic hernia (CDH) (Ijsselstijn, Gaillard et al. 1997, H, Hung et al. 1998). In addition, NE cells have proliferative potential and are the cell of origin driving development of human small cell lung cancer (van Meerbeeck, Fennell et al. 2011). The various diseases with a disturbed NE cell phenotype suggests that involvement of NE cells is of a complex nature.

Pulmonary NE cells are chemo-sensory cells, which monitor different aspects of lung physiology, such as changes in oxygen, other chemicals and changes in mechanical forces (Cutz, Pan et al. 2013). Changes registered by the NE cells are relayed to the brain by sensory neurons (Brouns, Oztay et al. 2009). In order to exert biological responses to the detected changes, NE cells contain dense core vesicles containing neuropeptides, such as Calcitonin gene-related peptide (CGRP) (Cutz, Pan et al. 2013). In the mouse lung, solitary NE cells are scattered throughout the airway epithelium, whereas clusters of NE cells, called neuroendocrine bodies (NEBs), are often found at airway bifurcations (Kuo and Krasnow 2015). In mouse models it was shown that upon allergen exposure, NE cells are activated, and initiate goblet cell hyperplasia and infiltration of immune cells into the lung (Sui, Wiesner et al. 2018). The increased mucus production due to goblet cell hyperplasia is initiated by the production and secretion of GABA by NE cells, which is completely abrogated upon loss of neuronal innervation (Barrios, Patel et al. 2017, Barrios, Kho et al. 2019). Besides the importance of NE cell innervation, a proper clustering of NE cells was shown to be essential in restricting immune infiltration in the neonatal lung (Branchfield, Nantie et al. 2016), showing a clear difference in functionality between solitary NE cells and NEBs.

NE cells are endoderm-derived, and together with basal cells are the earliest cells originating from the airway progenitors. Notch signaling plays an active role in distinguishing between a NE and non-NE cell fate (Ito, Udaka et al. 2000, Morimoto, Nishinakamura et al. 2012, Jia, Wildner et al. 2015). In non-NE cells, active Notch signaling represses the transcription of *Achaete-scute homolog 1* (*Ascl1*), which is a master regulator in the formation of NE cells (Borges, Linnoila et al. 1997). Previously, we showed that overexpression of the transcription factor sex determining region Y-box 2 (SOX2) in the developing lung epithelium resulted in increased expression of *Ascl1* and an increased number of NE cells (Gontan, de Munck et al. 2008). Moreover, we showed that early in development the expression levels of both SOX2 and SOX21 determine the fate of airway progenitor to basal cells (Eenjes, Buscop-van Kempen et al. 2021). In neural and olfactory precursor cells, ASCL1 has shown to be a transcriptional target of SOX2 (Tucker, Lehtinen et al. 2010, Niu, Zang et al. 2015). However, if and how SOX2 and SOX21 play a role in the initial differentiation and distribution of NE cells during lung development is not yet determined.

Here, we show that SOX2 and SOX21 have distinct roles in the development and maturation of NE cells. We show that reduced levels of SOX21, but not SOX2, affect the initiation of the formation of NE precursor cells. Both, SOX2 and SOX21 deficiency resulted in an aberrant clustering of NE cells compared to wild-type conditions. In addition, deficiency of SOX2 levels delayed NE cell maturation.

## RESULTS

### SOX21 inhibits SOX2 progenitor differentiation to precursor NE cells

In previous studies we suggested a relation between SOX2 and NE cell development (Gontan, de Munck et al. 2008). A potential interaction between SOX2 and SOX21 within the development of NE cells, was determined by evaluating whether NE precursors appear in the SOX2+SOX21+ proximal airway region (Eenjes, Buscop-van Kempen et al. 2021). At E13.5 and E14.5, solitary and clustered (~10-30 cells) NE precursor cells (ASCL1+) were observed in the proximal airways, consistent with previous published data (Kuo and Krasnow 2015, Noguchi, Sumiyama et al. 2015) (Figure 1A). We found that precursor NE cells, were observed in the region positive for SOX21, at both E13.5 and E14.5 (Figure 1B). We assessed SOX2 and SOX21 expression levels in ASCL1+ NE and ASCL-non-NE cells and calculated the mean fluorescence intensity (MFI) of SOX2 and SOX21 in ASCL+ and ASCL1-nuclei (Figure 1C). This showed no correlation between ASLC1+ precursor NE cells and MFI of SOX2 or SOX21 (Figure 1C). At E14.5, SOX2+ airway progenitor cells also differentiate to TRP63+ basal cells (Transformation Related Protein 63 positive cells). Similar to the precursor NE cells, we did not observe a correlation, between TRP63+ and TRP-nuclei and MFI of SOX2 or SOX21 at this stage of development (Figure 1D). Thus similar to basal cells, precursor NE cells start appearing in the airway regions positive for SOX2 and SOX21, but SOX21 or SOX2 abundancy does not correlate with either basal or NE cells early in lung development.

**Figure 1:**
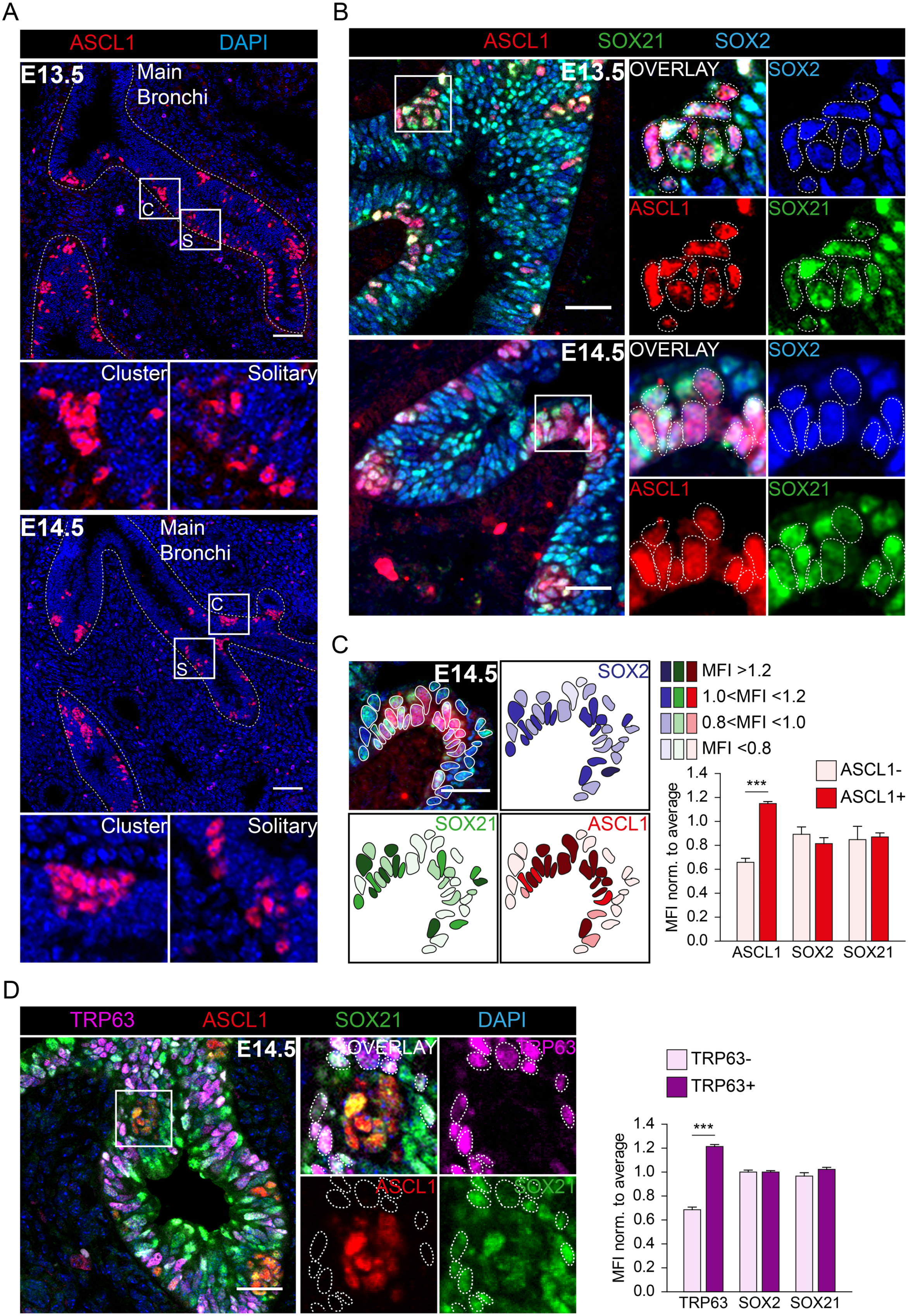
Differentiation to NE cells of SOX2+ airway progenitor cells occurs in SOX21+ airway region. (A) Immunofluorescence staining of ASCL1 (red) and DAPI (blue) at gestational ages E13.5 and E14.5. C=cluster, S=solitary. Scale bar = 50 μm. (B) Immunofluorescence staining of ASCL1 (red), SOX21 (green) and SOX2 (blue) at gestational age E13.5 and E14.5. Cells positive for ASCL1 are encircled. Scale bar = 25 μm. (C) Immunofluorescence staining of ASCL1 (red), SOX21 (green) and SOX2 (blue) at gestational age E14.5. In the encircled cells the MFI of ASCL1, SOX21 and SOX2 is shown at that specific region. Scale bar = 25 μm. The graph shows the MFI of ASCL1, SOX21 and SOX2 in ASCL- and ASCL+ airway epithelium. Data are represented as mean ± SD. Two-way ANOVA (n = 3, *** p<0.001). (D) Immunofluorescence staining of TRP63 (violet), ASCL1 (red), SOX21 (green) and DAPI (blue) at gestational age E14.5. Cells positive for TRP63 are encircled. Scale bar = 25 μm. The graph shows the MFI of TRP63, SOX21 and SOX2 in TRP63- and TRP63+ airway epithelium. Data are represented as mean ± SD. Two-way ANOVA (n = 3, *** p<0.001).

We next determined whether levels of SOX2 or SOX21 play a role in the differentiation of SOX2+ airway progenitors to NE cells. The number of precursor NE cells were counted at E14.5 in lungs of SOX21 heterozygous (*Sox21*^+/-^), knock-out (*Sox21^-/-^*) and SOX2 heterozygous (*Sox2*^+/-^) mice. Comparable numbers of precursor NE cells were observed in lungs of wild type (WT), *Sox21*^+/-^ and *Sox2*^+/-^. Although the numbers of NE cells were not significantly different in lungs of *Sox21^+/-^* mice, complete ablation of SOX21 resulted in increased numbers of precursor NE cells (Figure 2). So, absence of SOX21 increased the differentiation of SOX2+ airway progenitors to NE cells, indicating that SOX21 may control the emergence of NE cells. In addition, we observed that in both the *Sox21^+/-^* and *Sox21^-/-^* NE cells were more solitary distributed throughout the airway epithelium (Figure 2).

**Figure 2:**
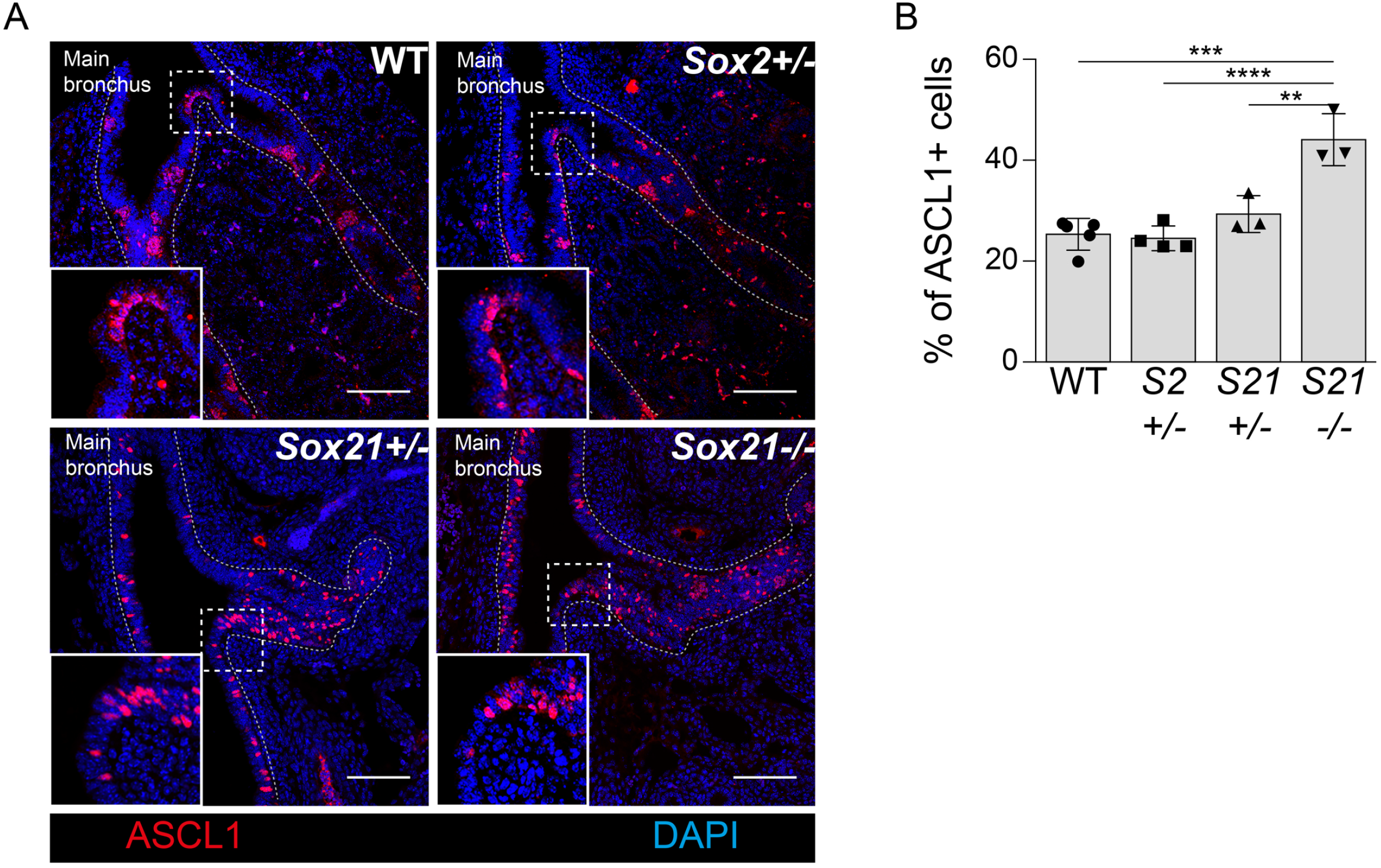
Loss of SOX21 increases differentiation of SOX2+ airway progenitor cell to NE precursor cells. (A) Immunofluorescence staining of ASCL1 (red) on the first bifurcation of the main bronchus at gestational age E14.5 of wild-type (WT), *Sox2^+/-^, Sox21^+/-^* and *Sox21^-/-^* mice. (B) Quantification of the percentage of NE precursor cells in a box of 400 μm^2^ around the first bifurcation of the main bronchus. Data are represented as mean ± SEM. One-way ANOVA (n WT = 5, n *Sox2^+/-^* = 4, *Sox21^+/-^* = 3, *Sox21^-/-^* = 3, ** p<0.01, *** p<0.001, ****p<0.0001).

### Expression levels of SOX21 decrease during the maturation of NE cells

To gain more insight in the role for SOX2 and SOX21 in the formation of NE cells during lung development, we looked at the development of NE cells at later gestational ages. Functional NE cells are innervated by sensory neurons and express the neuropeptide protein, CGRP.

Innervation of NE cells was determined by the presence of synaptic vesicle protein 2 (SV2), a general marker of synaptic vesicles in neuronal cells (Weichselbaum, Everett et al. 1996, Pan, Yeger et al. 2004). SV2 and CGRP were detected weakly at E16.5 and more abundantly expressed at E18.5 (Figure 3). From E15.5 onward, NE cells started to appear in the distal airways, probably due to migration of the NE cells in a proximal to distal fashion (Noguchi, Sumiyama et al. 2015). Distal airway epithelial cells express SOX2, but not SOX21, especially later in lung development (>E15.5) (Eenjes, Buscop-van Kempen et al. 2021). At E15.5, SOX21 was observed both in NEBs in the main bronchi and in the distal airways (Figure 4A, Figure 4-figure supplement 1). Resembling E14.5, in the main bronchi of E15.5, the MFI of SOX2 and SOX21 was similar in NE cells compared to the adjacent non-NE airway epithelium (Figure 4A). At E16.5, when the earliest expression of CGRP could be detected (Figure 3), NE cells in the main bronchi showed a lower MFI of SOX21 and a slightly lower MFI of SOX2 compared to adjacent non-NE airway epithelium (Figure 4A). Distal airway epithelium lacks SOX21, and we did not observe expression of SOX21 in NE cells at E16.5 (Figure 4-figure supplement 1)(Eenjes, Buscop-van Kempen et al. 2021). At E18.5, NEBs were both SOX2 and SOX21 positive in proximal and distal airway epithelium (Figure 4A, Figure 4-figure supplement 1). However, the MFI of SOX21 was slightly lower in NE cells compared to adjacent non-NE airway epithelium (Figure 4A). In conclusion, the levels of SOX2 and SOX21 are dynamically expressed during formation and differentiation of the NE cells. NE cells start to express CGRP around E16.5 and this is accompanied by a decrease in SOX21 levels. We also observed a small, but non-significant decrease in SOX2 levels at this developmental stage. At E18.5, SOX21 is observed again in NEBs (Figure 4B).

**Figure 3:**
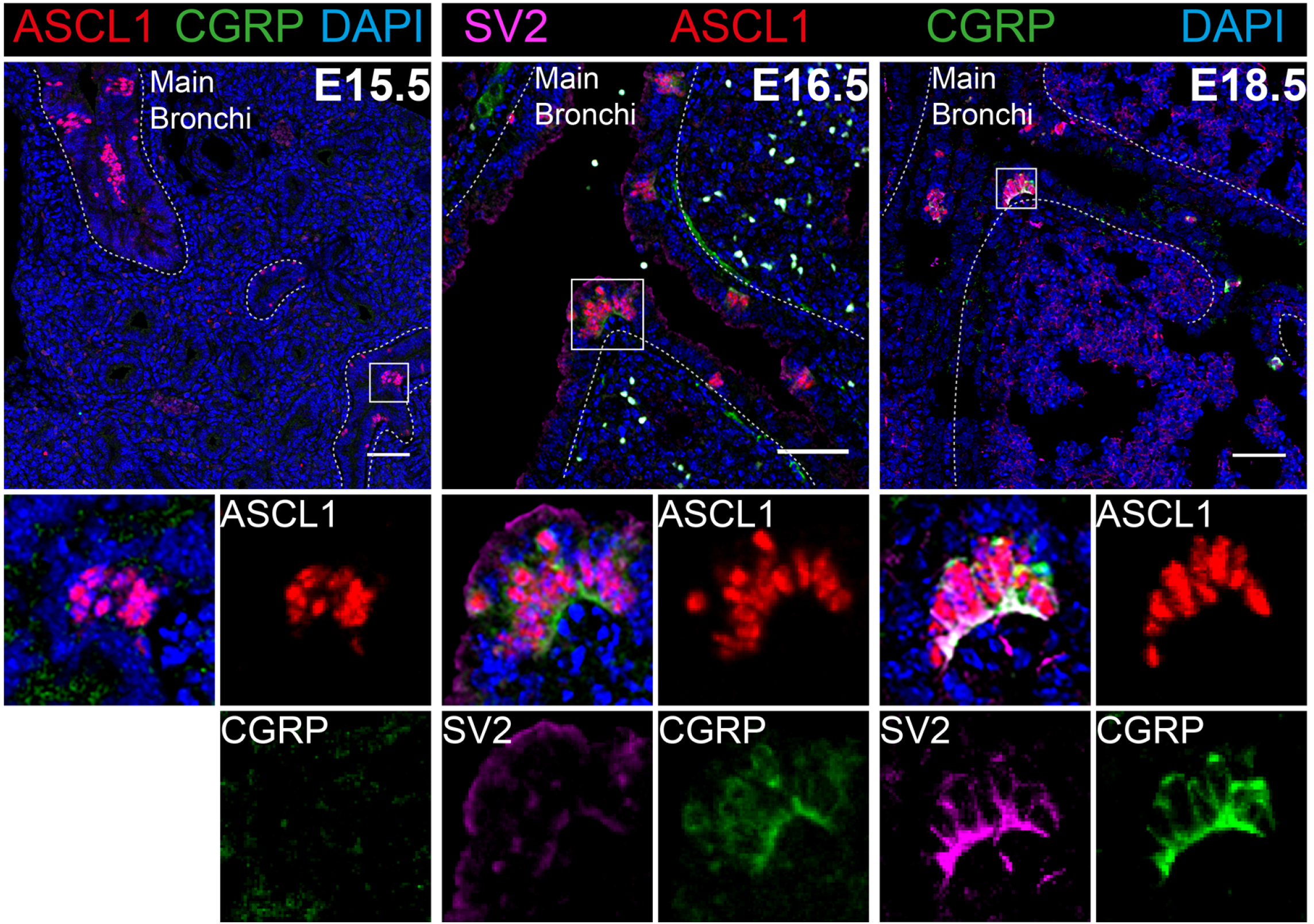
Innervation of NEBs and CGRP production starts at E16.5. Immunofluorescence staining of SV2 (violet), ASCL1 (red), CGRP (green) and DAPI (blue) at gestational ages E15.5, E16.5 and E18.5. Scale bar = 50 μm.

**Figure 4:**
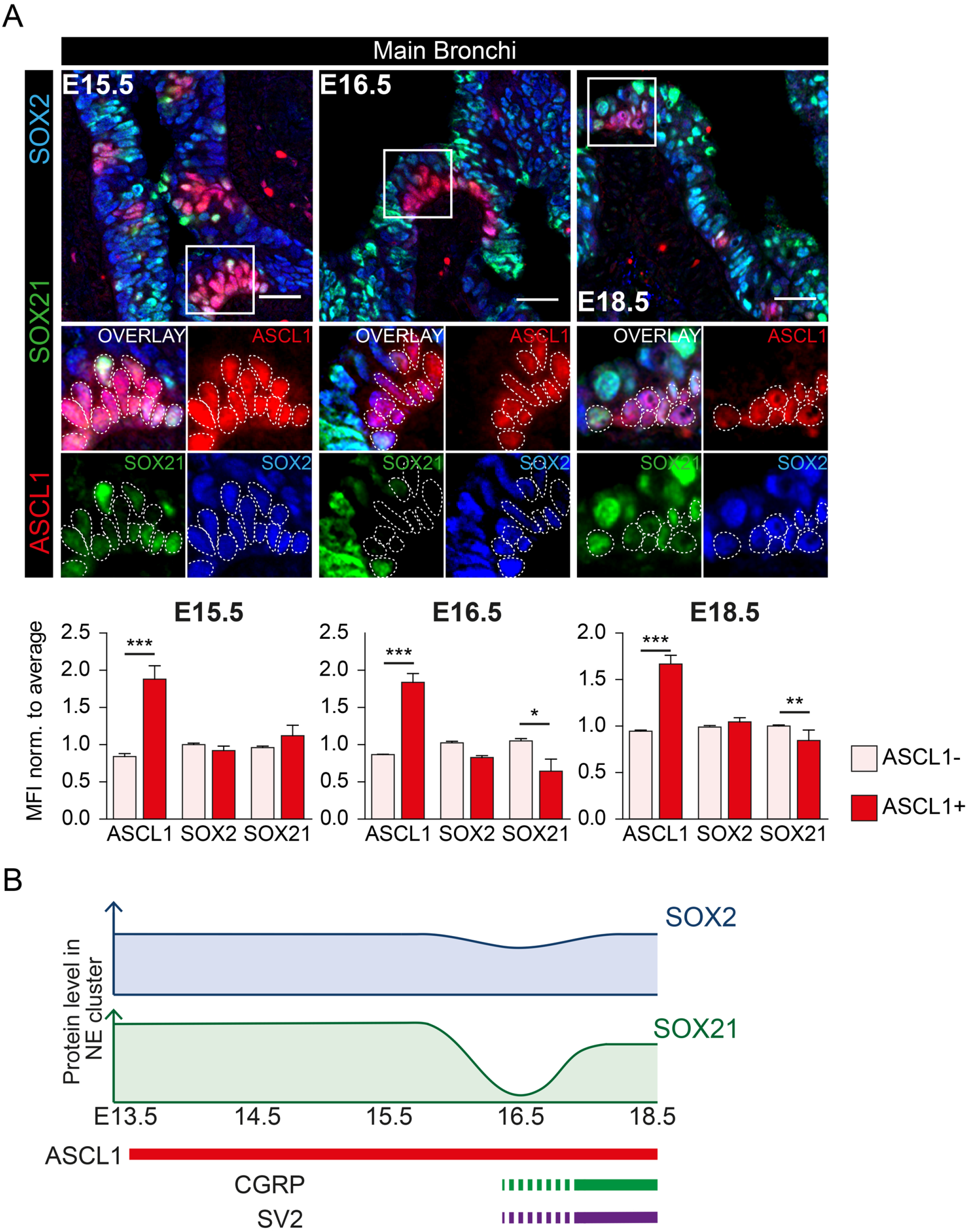
Different abundancy of SOX2 and SOX21 during the maturation of NEBs. (A) Immunofluorescence staining of ASCL1 (red), SOX21 (green) and SOX2 (blue) at gestational age E15.5, E16.5 and E18.5. The proximal airway epithelium is characterized by abundant expression of SOX21. Cells positive for ASCL1 are encircled. Scale bar = 25 μm. The graph shows the MFI of ASCL1, SOX21 and SOX2 in ASCL- and ASCL+ airway epithelium in proximal airway epithelium at E15.5, E16.5 and E18.5. Data are represented as mean ± SD. Two-way ANOVA (n = 3, * p<0.05, **p<0.01, *** p<0.001). (B) Schematic representation of SOX2 and SOX21 protein levels in neuroendocrine bodies (NEBs) at different gestational ages. SOX21 shows a dip and SOX2 a slight decrease in expression at E16.5, when CGRP expression starts and NEBs become innervated (SV2).

### SOX21 and SOX2 are important in the maturation and clustering of NE cells

Next, we analyzed the number of CGRP+ NE cells distributed throughout the E18.5 lung. Similar to E14.5, an increased number of NE cells was still present at E18.5 in the *Sox21^-/-^* lungs. Contrary to E14.5, where no difference in the number of NE precursors was detected, *Sox2^+/-^* E18.5 airways showed a decreased number of CGRP+ NE cells (Figure 5A, B). We observed that ASCL1+ NE cells at E18.5 in WT, *Sox21^+/-^* and *Sox21^-/-^* airways, were all mature NE cells (CGRP+), but the SOX2^+/-^ airways showed a substantial number of ASCL1+ NE cells that did not express CGRP (Figure 5C). In conclusion, SOX21 levels seem to suppress the initiation of NE cells early in lung development, where high SOX2 levels are important in the maturation of NE cells.

**Figure 5:**
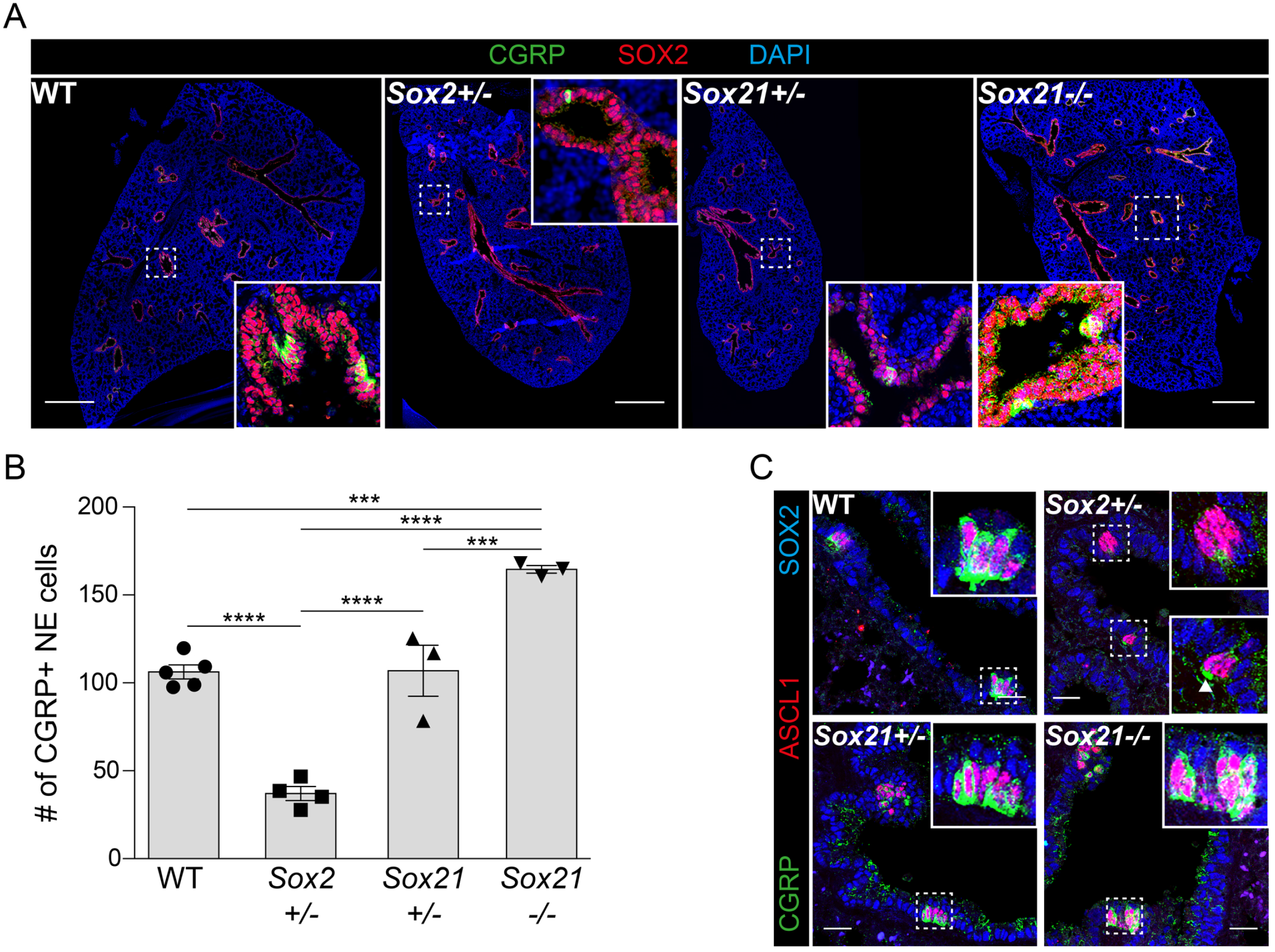
Loss of SOX2 decreases and of SOX21 increases the number NE cells at E18.5. (A) Immunofluorescence staining of CGRP (green), SOX2 (RED) and DAPI (blue) on lung sections at gestational age E18.5 of wild-type (WT), *Sox2^+/-^, Sox21^+/-^* and *Sox21^-/-^* mice. Scale bar = 500 μm. (B) Quantification of the number of NE cells on E18.5 lung sections, normalized to the area of SOX2+ area measured. Data are represented as mean ± SEM. One-way ANOVA (n WT = 5, n *Sox2^+/-^* = 4, *Sox21^+/-^* = 3, *Sox21^-/-^* = 3, ** p<0.01, *** p<0.001, ****p<0.0001). (C) Immunofluorescence staining of ASCL1 (red), CGRP (green) and SOX2 (blue) on lung sections at gestational age E18.5 of wild-type (WT), *Sox2^+/-^, Sox21^+/-^* and *Sox21^-/-^* mice. Scale bar = 25 μm.

In the mouse lung, NE cells are found solitary, in small clusters (2-5 PNECs) or big clusters / NEBs (>5 NE cells). The distribution and innervation of NE cells has been shown to be important for their function (Branchfield, Nantie et al. 2016, Barrios, Patel et al. 2017). We observed, that both clusters and single CGRP+ NE cells show neuronal innervation (SV2+) in WT, *Sox2^+/-^*, Sox21^+/-^ and *Sox21^-/-^* lungs (Figure 6A). Next, we determined the distribution of NE cells upon reduced levels of *Sox2* and *Sox21* expression (Figure 6A). In WT lung, NE cells were mostly present in small and big clusters, while in *Sox2^+/-^* lungs, NE cells were slightly more present as solitary cells although most of the NE cells were present in small clusters (Figure 6B). Reduced expression levels or complete absence of SOX21, resulted both in a high percentage of NE cells in NEBs and smaller percentage as solitary NE cell (Figure 6B). All genotypes contained a similar number of cells in NEBs (Figure 6C). Taken together, the data show that SOX21 levels play a role in both initiation of NE cell differentiation and the distribution of cells in NEBs, while SOX2 levels are involved in the maturation of NE cells and the clustering of NE cells in NEBs.

**Figure 6:**
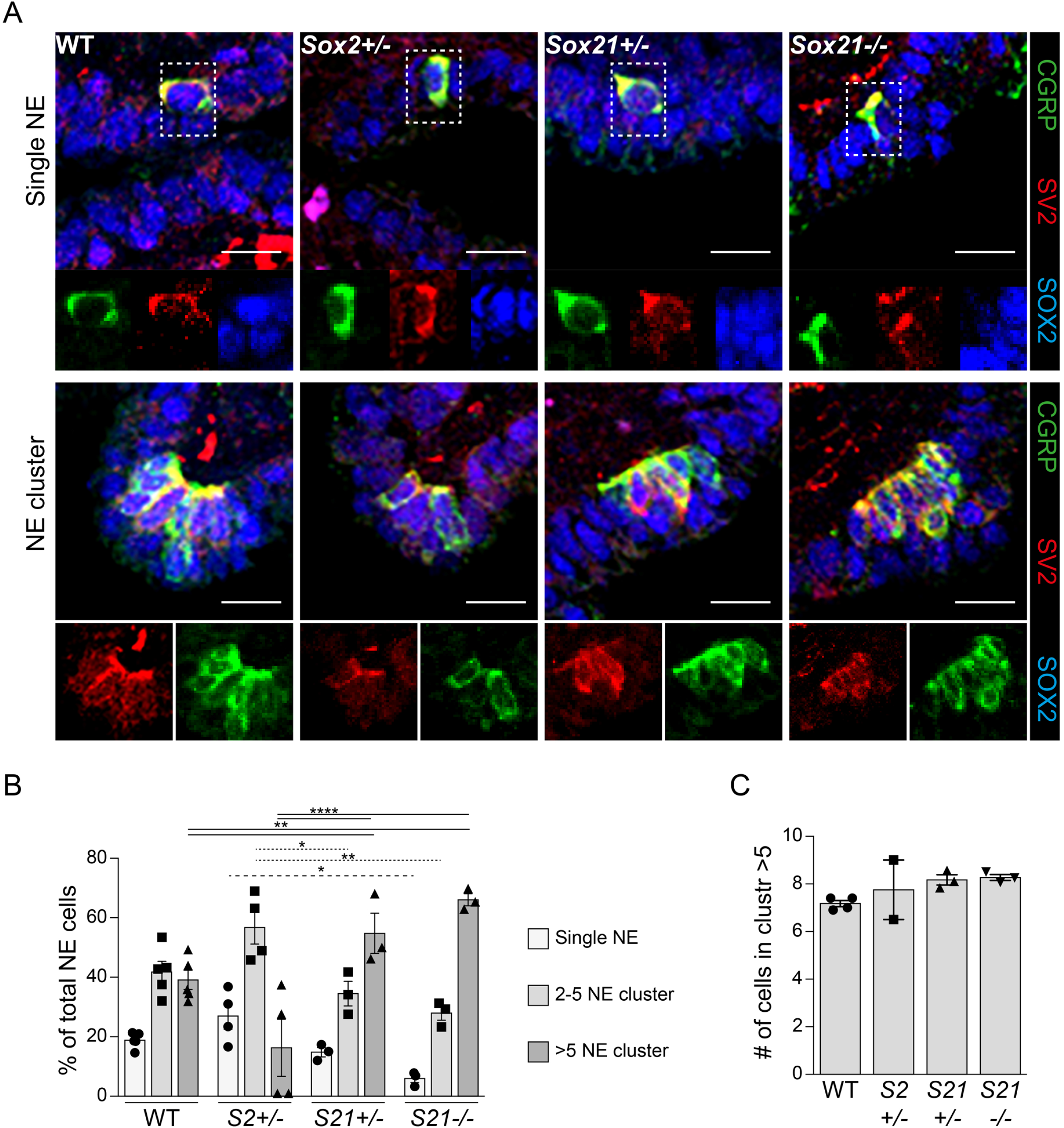
Loss of SOX21 increases the number NE clusters. (A) Immunofluorescence staining of CGRP (green) and SV2 (RED) on lung sections at gestational age E18.5 of wild-type (WT), *Sox2^+/-^, Sox21^+/-^* and *Sox21^-/-^* mice. Scale bar = 15 μm. (B) The percentage of NE cells that are present as single NE cells, small clusters (2-5 cells) or large clusters (>5). Data are represented as mean ± SEM. Two-way ANOVA (n WT = 5, n *Sox2^+/-^* = 4, *Sox21^+/-^* = 3, *Sox21^-/-^* = 3, * p<0.05, ** p<0.01, ****p<0.0001). (C) Representation of the average number of NE cells in clusters >5. One-way ANOVA (n WT = 5, n *Sox2^+/-^* = 4 (2 did not show any clusters >5), *Sox21^+/-^* = 3, *Sox21^-/-^* = 3)

## DISCUSSION

Both SOX21 and SOX2 were known to be co-expressed in developing proximal airway epithelium (Eenjes, Buscop-van Kempen et al. 2021), but their individual functions in the development of NE cells were not known. In this study, we investigated their potential involvement in the initiation, maturation and distribution of NE cells in the mouse lung. Deficiency in either SOX2 or SOX21 expression, shows that SOX21 suppresses the initial differentiation to NE cells early in development while SOX2 levels are mainly important in the maturation of NE cells later during lung development.

During airway development, airway progenitor cells differentiate to basal, secretory, ciliated and NE cells. Complete deficiency of the Notch signaling pathway, through deletion of *Rbpjk* (Notch transcriptional effector), *Pofut1* (essential for Notch-ligand binding) (Tsao, Vasconcelos et al. 2009) or deletion of all Notch receptors (Morimoto, Nishinakamura et al. 2012), resulted in the expansion of NE cells. The initial stages of airway progenitor differentiation to NE cells, takes place in the SOX2+SOX21+ intrapulmonary airways. Here, we show that SOX21 is important in suppressing NE cell differentiation, which is dependent on the level of expression as demonstrated by SOX21 hetero- and homozygous knockout mice.

Based on our data we hypothesize that a competition between SOX and SOX21 exists and that subtle changes in the balance between these factors determines cell fate decisions. This is supported by the physically interaction between SOX2 and SOX21 (Mallanna, Ormsbee et al. 2010, Kuzmichev, Kim et al. 2012), by the transcriptional activation of SOX21 by SOX2 (Gontan, de Munck et al. 2008, Kuzmichev, Kim et al. 2012), and by the facts that SOX2 and SOX21 bind the same genomic locations, and SOX21 can compete SOX2 binding to suppress gene activation (Eenjes, Buscop-van Kempen et al. 2021). Together, these data support the hypothesis that SOX2 and SOX21 compete for DNA binding sites, and thus changes in the balance between SOX2 or SOX21 directly influence the binding of these factors to DNA. As a result, targets genes are either activated, or not, because SOX2 has a C-terminal transactivation domain, whereas SOX21 harbors a transcriptional repressor motif in the C-terminal part of the protein (Kamachi and Kondoh 2013).

In addition, *Sox2^+/-^* mice did not show an increase or decrease in the number of NE precursor cells, showing that deficiency in SOX2 or SOX21 does not affect the initial differentiation to NE cells. In neuronal stem cell differentiation, SOX21 deletion also increased differentiation to ASCL1+ precursor cells through suppression of Notch target HES5 (Matsuda, Kuwako et al. 2012). NE cell differentiation in the developing lung is repressed by HES1 in non-NE airway epithelial cells(Ito, Udaka et al. 2000, Jia, Wildner et al. 2015).

Although, we did not observe a difference in the number of precursor NE cells upon reduced levels of SOX2, we did show that SOX2 plays an important role in the expression of CGRP in NE cells at the end of lung development. These results indicate that high levels of SOX2 are not necessary for the induction of *Ascl1* but are necessary for maturation of NE cells. Furthermore, reduced levels of SOX2 showed a higher frequency of single scattered NE cells in the airway epithelium, while reduced levels of SOX21 resulted in less solitary NE cells but more NEBs. A difference in abundancy of SOX2 and SOX21 were observed within NEBs at different stages of NE development. This, all together suggests that SOX2 and SOX21 might function differently during different stages of NE development; initiation, migration and maturation.

In addition to the function of the NE cell as a chemo-sensory cell, a subset of NE cells was shown to proliferate and repair surrounding airway epithelium upon response to injury (Reynolds, Hong et al. 2000, Ouadah, Rojas et al. 2019). In several mouse models regulation of maintenance and proliferation of NE cells is important, because NE cells were found to be the origin for small cell lung cancer (SCLC) (Sutherland, Proost et al. 2011, Song, Yao et al. 2012). It has long been speculated that SCLC arise from NE cells due to similarities in morphology and expression of NE markers in the tumor (van Meerbeeck, Fennell et al. 2011). It was recently established that specific NE stem cells are the source of tumor formation within NEBs (Ouadah, Rojas et al. 2019). Multiple studies have shown an increase in *Sox2* expression in SCLC, and a few also indicated elevated levels of *Sox21* (Gure, Stockert et al. 2000, Titulaer, Klooster et al. 2009, Rudin, Durinck et al. 2012, Zhu, Li et al. 2012).

In conclusion, this study shows distinct involvement of SOX2 and SOX21 in the development and maturation of pulmonary NE cells. SOX21 suppresses initiation of NE cell differentiation early in development, preventing NE cell hyperplasia within the lung. In contrast, high SOX2 levels are mostly involved in the maturation and clustering of NE cells. Given that hyperplasia of NE cells and defective clustering are associated with several lung diseases and NE function respectively, investigating the molecular mechanisms downstream of SOX2 and SOX21 may provide new insight in NE function and development of new therapeutic approaches.

## MATERIAL AND METHODS

### Mice

All animal experimental protocols were approved by the animal welfare committee of the veterinary authorities of the Erasmus Medical Center. Mice were kept under standard conditions. Mouse strains *bioSOX2flox* (Schilders, Eenjes et al. 2018) and *SOX21-KO* (gift of Stavros Malas) were used. Wild-type animals were C57BL/6.

### Immunofluorescence

Mouse embryonic lungs were fixed overnight in 4% w/v PFA at 4 °C. Post-fixation, samples were washed with PBS, de-hydrated to 100% ethanol, transferred to xylene and processed to paraffin wax for embedding. Paraffin blocks were sectioned at 5 μm and dried overnight at 37 °C.

Sections were deparaffinized by 3 times 2 min xylene washes, followed by rehydration in distilled water. Antigen retrieval was performed by boiling the slides in Tris-EDTA (10mM Tris, 1mM EDTA) buffer pH=9.0 for 15 min at 600W. Slides were cooled down for 30 min and transferred to PBS. For SOX21 and ASCL1 staining, the Tyramide Signal Amplification (TSA) kit was used (Invitrogen, B40922, according to manufacturer’s protocol). When using the TSA kit, a hydrogen peroxide (1.5% in PBS) blocking step was performed after boiling. For co-staining of SOX21 and ASCL1, first ASCL1 was stained (first and secondary antibodies plus TSA reaction), sections were boiled again in Tris-EDTA (10mM Tris, 1mM EDTA) buffer pH=9.0 for 15 min at 300W and subsequently SOX21 was stained. Sections were blocked for 1 hr at room temperature (RT) in 5% Elk (Campina) for ASCL1 staining or 3% BSA, 0.05% Tween in PBS. Primary antibodies (table 1) were diluted in blocking buffer and incubated with the sections overnight at 4 °C. The next day, sections were washed 3 times for 5 min at room temperature (RT) in PBS. Secondary antibodies (table 1) were added in blocking buffer and incubated for 1 hr at RT. DAPI (4’,6-Diamidino-2-Phenylindole) solution (BD Pharmingen, 564907, 1:2000) was added to the secondary antibodies for nuclear staining. After incubation, 3 times 5 min washes in PBS, sections were mounted using Mowiol reagent (For 100 mL: 2,4% w/v Mowiol, 4,75% w/v glycerol, 12 % v/v Tris 0.2M pH=8.5 in dH_2_O till 100 mL). All sections were imaged on a Leica SP5 confocal microscope.

**Table 1.**
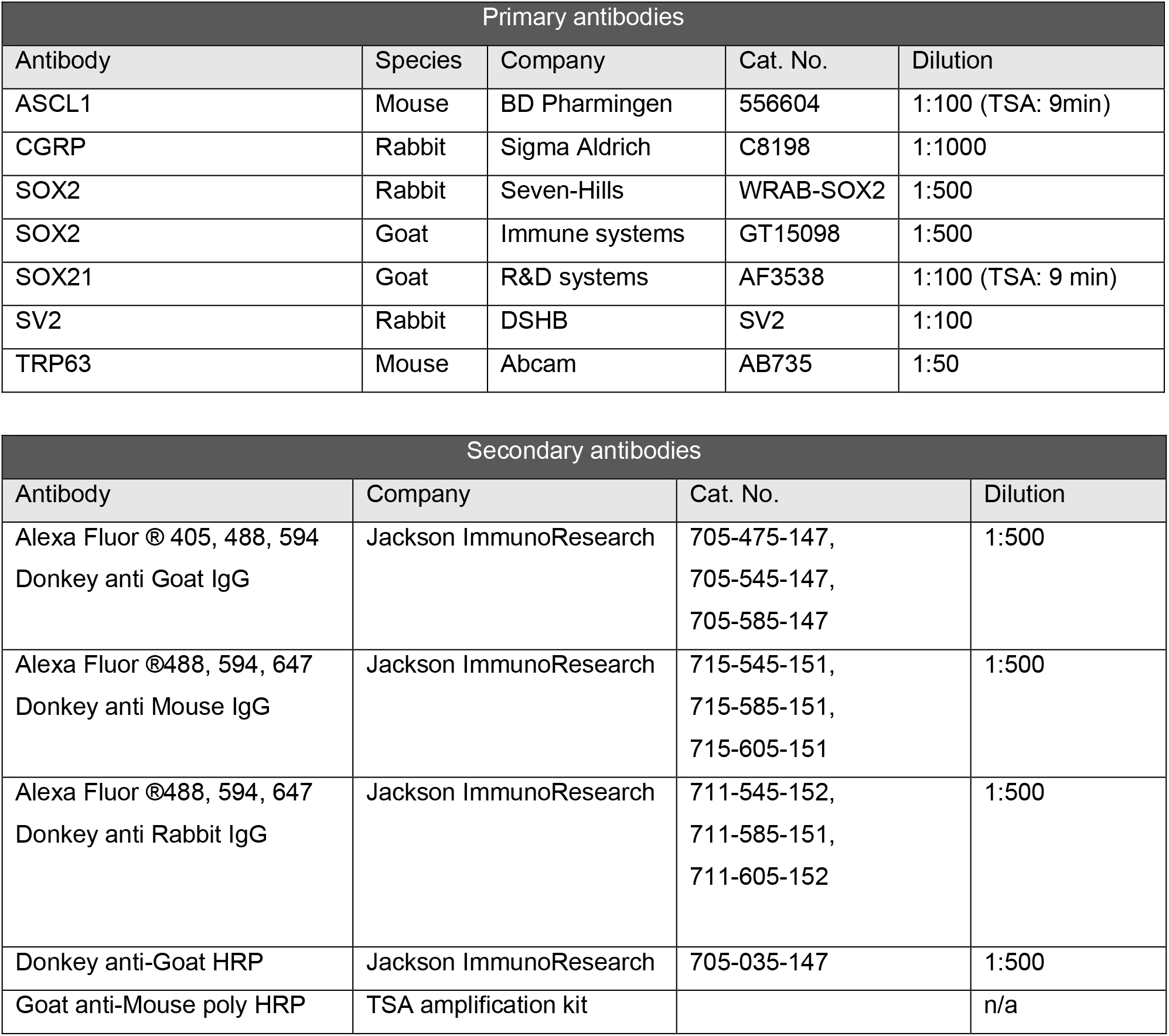
Antibodies

### Image analysis

Neuroendocrine precursor cells (ASCL1+) were counted on E14.5, 5 μm thick sections, in a square of 400 μm^2^ around the first branch of the main bronchi. Of each genotype and each n, 3 sections were counted and the percentage of NE cells were calculated based on the total number of DAPI+ nuclei in the airway.

Neuroendocrine cells (CGRP+) were counted on E18.5, 5 μm thick sections. Of each genotype and each n, 2 complete sections were counted and the number of NE cells were normalized to airway (SOX2+) area present in the section.

Fluorescent intensity of SOX2 and SOX21 was measured in three different location which contained ASCL1+ NEBs, in proximal SOX2+SOX21+ airway epithelium. The mean fluorescence intensity (MFI) was calculated by dividing the fluorescence intensity of ASCL1+ or ASLC1-region compared to the fluorescence intensity of the total airway epithelium at that specific location.

Fluorescent intensity of TRP63, SOX2 and SOX21 was measured in a 150 μm^2^ box in the main bronchi, in 3 sections of 3 different mouse lungs. The MFI was calculated by dividing the fluorescence intensity of TRP63+ or TRP63-region compared to the fluorescence intensity of the total airway epithelium at that specific location.

Image J was used to analyze the pictures.

## ACKNOWLEDGEMENTS

We like to thank Frank Grosveld and Niels Galjart for critically reading the manuscript (Department of Cell Biology, Erasmus MC). This work was supported by grants from the Sophia Foundation for Medical Research S14-12 (EE).

---

**Figure 4-figure supplement 1:**
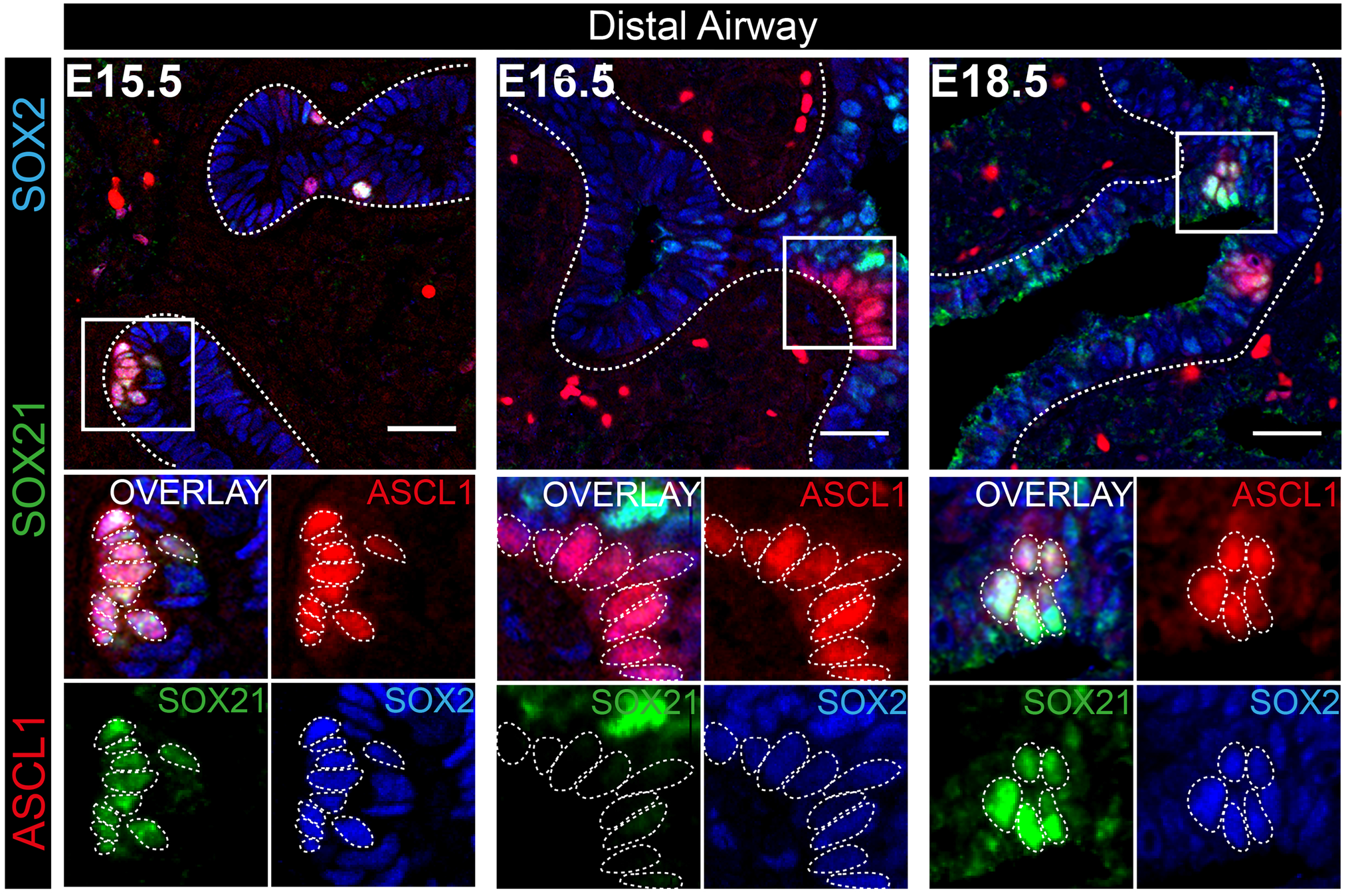
Expression of SOX2 and SOX21 during the maturation of NEBs in the distal airway epithelium. Immunofluorescence staining of ASCL1 (red), SOX21 (green) and SOX2 (blue) at gestational age E15.5, E16.5 and E18.5 in the distal SOX2+ SOX21-airway epithelium. Cells positive for ASCL1 are encircled. Scale bar = 25 μm.

